# Saturation mutagenesis map of generalist versus specialist adaptations of β-lactamase to novel antibiotics

**DOI:** 10.1101/2025.10.14.682469

**Authors:** Ilona K. Gaszek, Muhammed S. Yildiz, Levent Sari, Ayesha Ahmed, Erdal Toprak, Milo M. Lin

## Abstract

The evolution of β-lactamase proteins is shaped by the need to maintain enzymatic activity against previously prevalent β-lactam antibiotics while expanding substrate range against new classes of antibiotics. Using saturation mutagenesis and sequence-barcoding-based quantification, we comprehensively mapped the response of the fitness landscape of TEM-1 β-lactamase, which evolved against penicillin-class antibiotics, to mutational perturbations against six diverse β-lactam antibiotics. This systematic panel of antibiotic substrates, including representatives from penicillin, cephalosporin, and monobactam classes, allowed us to classify resistance mutations into two categories. Generalist mutations conferred resistance to multiple antibiotics and were consistently restricted to three positions critical for substrate recognition and catalytic function R164, G238, and E240. These substitutions produced broad spectrum resistance through mechanisms such as expansion of the active site and improved substrate accommodation. In contrast, specialist mutations conferred resistance to only a single antibiotic and exhibited much wider positional diversity. Ceftazidime selection yielded the greatest number of distinct specialist mutations, which were frequently found in flexible or peripheral regions including the omega loop. One especially unexpected finding was the identification of the E166P variant. E166 is a catalytic residue required for deacylation during hydrolysis, and substitutions at this site are generally assumed to abolish function. However, E166P conferred a significant increase in ceftazidime resistance despite eliminating activity against penicillins. Molecular dynamics simulations and mutational analysis revealed that the E166P mutant employs an alternative catalytic mechanism, involving residue N132, rather than the canonical pathway. Together, our findings reveal, at the molecular level, how specialist mutations open up a wide range of diverse and idiosyncratic solutions at the expense of generalizability. These insights may inform strategic design of antibiotic administration protocols to systematically lower pathogenic evolvability.

## Introduction

The alarming rise in β-lactam antibiotic resistance, especially among Enterobacteriaceae, underscores the urgent need to understand the evolutionary flexibility of β-lactamase enzymes such as TEM-1 [1–3]. Originally optimized for penicillin hydrolysis and early generations of cephalosporins, TEM-1 has extended its substrate spectrum through the accumulation of point mutations, enabling resistance to a broader array of β-lactam antibiotics, including newer generations of cephalosporins and monobactams [4–6]. While numerous resistance-conferring mutations have been described, most previous studies have examined them in the context of one or two antibiotics, without systematically exploring how single-point mutations perform across diverse β-lactam classes [7, 8]. As an enzyme on the cusp of evolving specificity to a range of novel substrates, TEM-1 is a medically urgent case study for elucidating the molecular principles behind enzyme specificity versus broad activity. Such principles would not only inform our incomplete understanding of the fundamental constraints of enzyme optimization but would also be crucial for predicting evolutionary trajectories and informing therapeutic strategies combating β-lactam resistance.

To identify single-point mutations that initiate and promote the evolution of antibiotic resistance in TEM-1 β-lactamase, we employed a comprehensive saturation mutagenesis library [8] and tested this library via high-throughput resistance profiling under selective pressure from β-lactam antibiotics spanning penicillins, cephalosporins, monobactams, and carbapenems. We introduce a conceptual framework distinguishing between generalist positions, which can be mutated to confer broad-spectrum resistance to multiple β-lactams and specialist positions, which can only be mutated to confer resistance to a specific drug.

We show that there are only three generalist positions located at the active site, whereas over a dozen specialist mutations included both positions close to and distant from the active site. Ceftazidime-specific specialist mutations are especially numerous, many of which clustered around the Ω-loop, a structurally flexible region involved in substrate recognition. An unexpected discovery was the identification of E166P, which conferred a specific resistance only to ceftazidime despite abolishing native resistance to penicillins. In the canonical catalytic pathway, E166 activates the catalytic water molecule required for deacylation, a step crucial for hydrolyzing the β-lactam and regenerating the free enzyme [9, 10]. As such, mutations at E166 have long been considered evolutionarily constrained [11, 12]. We show that E166P confers activity against cetazidime via an alternative pathway involving N132 and modified water coordination.

Our investigation offers new insight into how point mutations at both structural and catalytic residues contribute to resistance, and how specialist adaptations like E166P may represent non-canonical evolutionary strategies that bypass established mechanistic constraints. Understanding the distinction and interplay between generalist and specialist mutations is crucial for predicting evolutionary trajectories and informing therapeutic strategies aimed at combating β-lactam resistance.

## Results

### Selection of Antibiotic-Resistant Variants from the TEM-1 Single Mutant Library

Understanding the molecular basis of antibiotic resistance requires a comprehensive mapping of mutational effects across entire proteins. Here, we investigated TEM-1 β-lactamase using a saturation mutagenesis library where every position of the mature protein was substituted with all possible amino acids. To mimic environmental conditions, a Single Mutant Library (SML) of TEM-1 β-lactamase was constructed as described previously by Stiffler et al. [8] on the low-copy plasmid pBR322. To identify antibiotics against which the SML exhibits increased resistance, we determined Minimal Inhibitory Concentrations of the library against representatives from four major β-lactam classes: penicillins (ampicillin), cephalosporins (cefotaxime, ceftriaxone, ceftazidime, cefepime), monobactams (aztreonam), and carbapenems (meropenem, imipenem) (**Supplementary Figure 1**). Also, we did not observe increase in resistance toward any drugs from the carbapenem group.

To gain information about key point mutations that increase resistance toward tested antibiotics, we developed a high-throughput sequencing-based assay to evaluate resistance profiles against the six β-lactam antibiotics that showed potential for resistance development. This assay allowed us to simultaneously challenge the pooled library with various concentrations of antibiotics in 96-well plates, with experiments performed in triplicate and growth monitored continuously over 18 hours. Wild-type TEM-1 was exposed to the same range of antibiotic concentrations in parallel with the SML under identical conditions. Looking at the growth patterns of wild-type TEM-1 across different antibiotic concentrations, we established selection thresholds where the wild-type enzyme no longer conferred protection, but mutant variants with enhanced catalytic activity might still enable bacterial survival. This selection created an evolutionary bottleneck that allowed only the most resistant TEM-1 variants to emerge from the SML population. For further analysis, we selected the following concentrations: ampicillin (4096 µg/ml) representing penicillins; ceftazidime (1µg/ml), cefotaxime (0.25µg/ml), ceftriaxone (0.25µg/ml), and cefepime (1µg/ml) representing cephalosporins; and aztreonam (0.5µg/ml) representing monobactams (**Supplement Figure 2**). Cultures exhibiting growth at these selection concentrations were transferred to 20 ml of antibiotic-free growth media to allow the surviving mutants to increase cell density. This step was essential to achieve sufficient plasmid concentrations for subsequent isolation and Illumina sequencing analysis (**Figure 1**). This approach enabled us to comprehensively identify the specific mutations conferring resistance to each antibiotic class.

**Figure 1.**
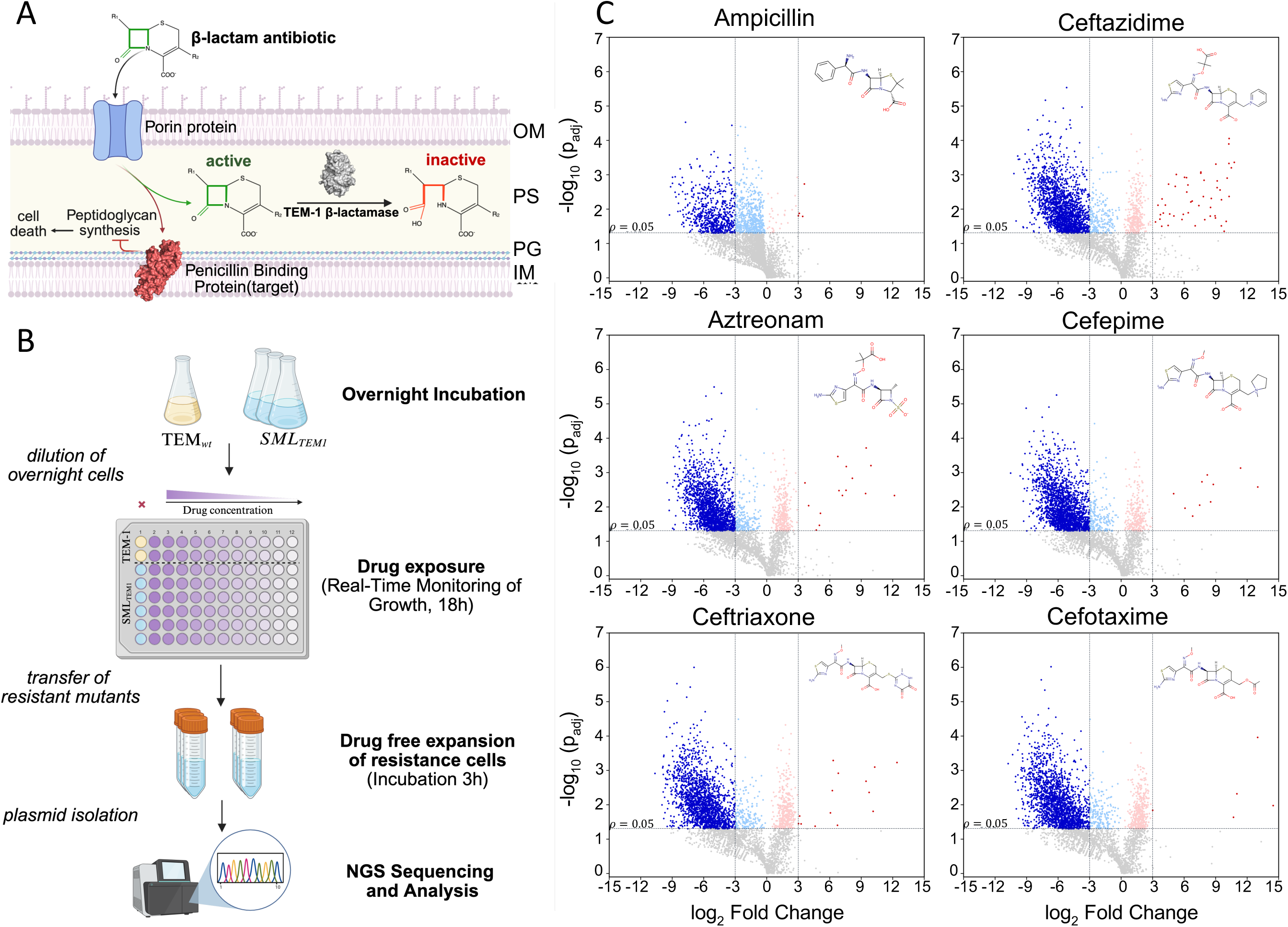
Saturation Mutagenesis of TEM-1 β-Lactamase Under diverse β-Lactam Antibiotic Pressure. **(A)** Mechanism of β-lactam antibiotic action and β-lactamase resistance. β-lactam antibiotics target bacterial cell wall synthesis, leading to cell death in susceptible bacteria. Bacteria producing β-lactamase enzymes hydrolyze the antibiotic, conferring resistance. OM-Outer Membrane, IM-Inner Membrane, PS-Periplasmic Space, PG-Peptidoglycan **(B)** Schematic overview of the experimental pipeline used to quantify fitness effects of TEM-1 β-lactamase variants in response to various β-lactam antibiotics. **(C)** Volcano plots obtained after selection experiments, highlighting differential fitness effects across conditions.

### High-throughput sequencing reveals distinct patterns of resistance-conferring mutations

To identify specific mutations conferring resistance to each β-lactam antibiotic, we isolated plasmids from surviving cultures and performed Illumina sequencing with 150bp paired-end reads. Following Illumina paired-end sequencing (150bp), we processed the data to quantify the abundance of each TEM-1 variant within each sample. Read counts were normalized to account for differences in sequencing depth across samples. To calculate mutant fitness, we implemented a robust statistical framework that compares the enrichment or depletion of each variant relative to its abundance in untreated conditions. Specifically, the fitness of each mutant variant was calculated as the ratio of its normalized abundance in treated versus untreated samples. For statistical analysis, we used a global synonymous mutation baseline derived from the median fitness of all synonymous mutations across the dataset to control for experimental variability. For each mutation, we computed the mean fold change relative to the synonymous baseline and performed statistical analysis using paired t-test on log₁₀-transformed values to evaluate the significance of observed fitness effects. The final fold changes were converted to log₂ scale for visualization. P-values were adjusted for multiple hypothesis testing using the Benjamini-Hochberg false discovery rate (FDR) correction with a significance threshold of 0.05. Significant mutations were visualized on volcano plots displaying log₂FoldChange on the x-axis and -log₁₀ (p-value) on the y-axis (**Figure 1C**). Mutations were classified as significantly beneficial when they exhibited both a log₂FoldChange > 3 (representing approximately 8-fold increased fitness compared to wild-type) and an adjusted p-value < 0.01. This stringent threshold ensured that we identified only the most robust resistance-conferring mutations. To further validate our analysis, we applied a secondary significance threshold (p < 0.05) to identify mutations with more modest but still statistically significant effects. Under these stringent criteria, only a small subset of mutations enabled TEM-1 variants to outperform the wild-type enzyme against each antibiotic. As expected, most mutations were either deleterious or had negligible impact on fitness, consistent with TEM-1’s evolutionary optimization for ampicillin hydrolysis.

Interestingly, ceftazidime selection yielded the highest number of beneficial mutations, correlating with the largest observed difference in MIC between resistant variants and wild-type TEM-1 compared to other tested β-lactams. This suggests that TEM-1’s structure presents multiple accessible evolutionary pathways toward ceftazidime resistance, potentially explaining the frequent emergence of ceftazidime-resistant variants in clinical settings.

### Classification of specialist and generalist resistance mutations

To further characterize the evolutionary landscape of antibiotic resistance, we classified beneficial mutations (log₂FoldChange > 3, padj < 0.05) into two categories: “specialist” mutations, conferring resistance to only one antibiotic, and “generalist” mutations, providing cross-resistance to two or more tested antibiotics. This classification highlights distinct evolutionary pathways leading to either narrow- or broad-spectrum resistance.

Generalist mutations are more predictable, localizing exclusively to three critical positions: R164, G238, and E240 (**Figure 2A, 2C**). These residues occupy strategically essential locations that directly control core aspects of catalytic function. G238 and E240 are situated at the edge of the active site, governing substrate binding specificity and interactions with β-lactam substrates. R164 is located within the critical Ω-loop, where it forms an essential salt bridge with D179 that stabilizes the catalytic residue E166 (also localized on the Ω-loop). This R164-D179 salt bridge ensures optimal hydrolysis rates for ampicillin but constrains accommodation of later-generation cephalosporins[13]. Mutations at R164 disrupt this salt bridge, increasing Ω-loop flexibility and enabling better accommodation of bulkier cephalosporins with improved E166 positioning for their deacylation [14, 15].

**Figure 2.**
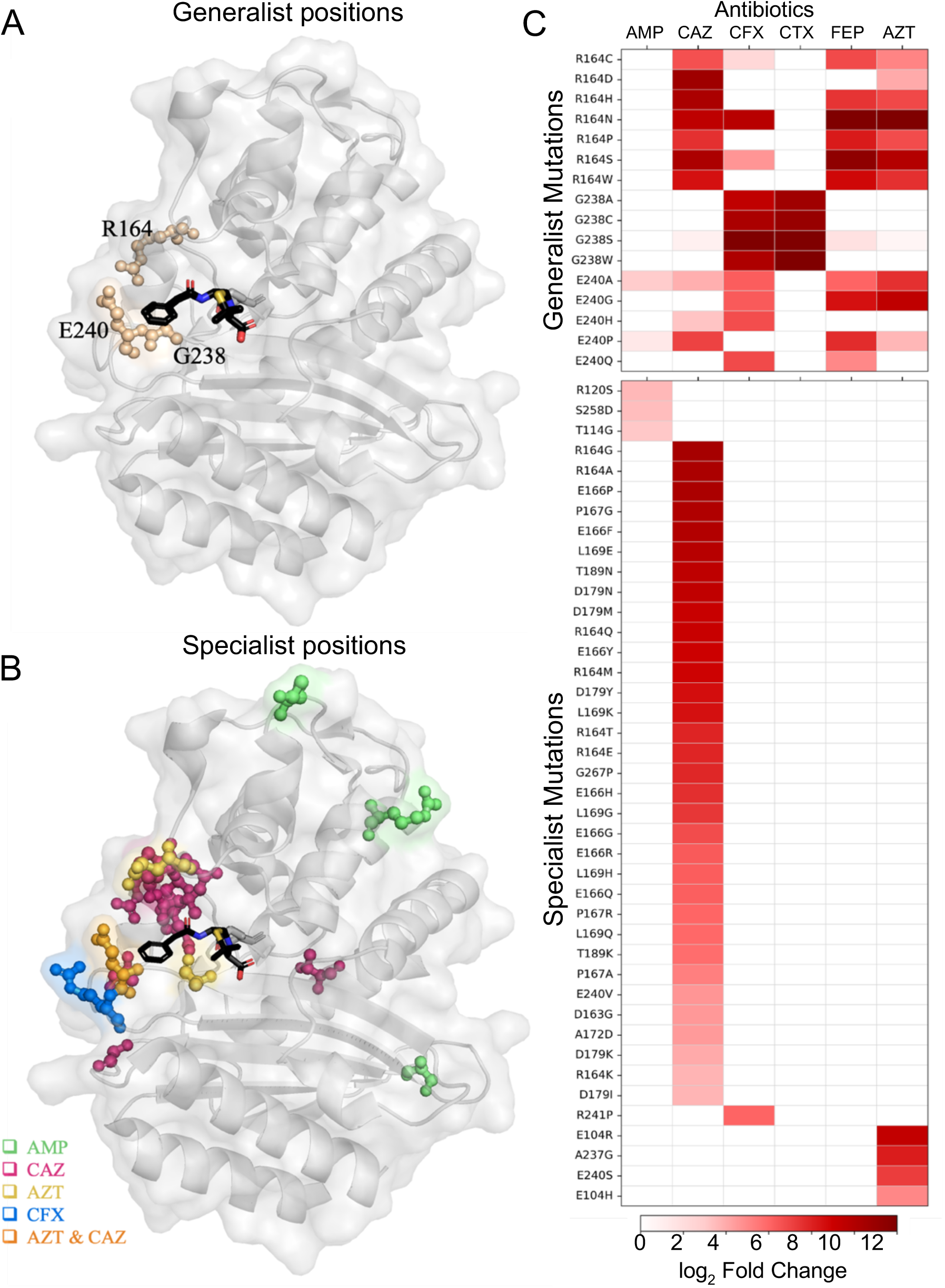
Structural Distribution and Classification of Specialist and Generalist Resistance Mutations in TEM-1 β-Lactamase. **(A)** Structural localization of generalist mutation positions on the TEM-1 β-lactamase structure. Generalist positions (R164, G238, and E240) are shown as wheat-colored spheres and are predominantly localized within or adjacent to the active site cavity. **(B)** Structural distribution of specialist mutation positions on the TEM-1 β-lactamase structure. Specialist positions are color-coded by antibiotic specificity: ampicillin specialists (AMP) green, ceftazidime specialists (CAZ) magenta, ceftriaxone specialists (CFX) blue, aztreonam specialists (AZT) yellow, and dual aztreonam/ceftazidime specialists are colored in orange. No significant specialist mutations were observed for Ceftriaxone (CTX) and Cefepime (FEP). **(C)** Resistance profiles of generalist and specialist mutations across β-lactam antibiotics. Two heatmaps displaying log₂ fold change values for TEM-1 mutations across six β-lactam antibiotics (AMP, CAZ, CFX, CTX, FEP, AZT). Top panel: Generalist mutations showing mutations that confer significant resistance (log₂ fold change > 3, padj < 0.05) to at least two antibiotics. Color intensity indicates the magnitude of fitness enhancement. Bottom panel: Specialist mutations showing mutations that confer significant resistance (log₂ fold change > 3, padj < 0.05) to exactly one antibiotic, demonstrating narrow-spectrum resistance profiles

Mutations at these strategic positions employ distinct molecular mechanisms. E240 variants stabilize interactions with the larger cephalosporin side chains [16], while G238S and R164 substitutions increase active-site flexibility through controlled destabilization [13, 17–20].

Our comprehensive saturation mutagenesis approach identified a broader mutational landscape of generalist mutations at these crucial sites compared to those typically observed clinically. Established clinical variants such as G238S and R164S were confirmed, but numerous additional potent generalist substitutions at these same positions, such as E240Q, E240H, G238C, and G238A were identified experimentally but are rarely or never observed clinically [21]. Conversely, the E240K mutation, commonly found in clinical isolates, provides limited fitness benefits on its own and typically requires additional mutations for effective resistance [5] .

An illustrative example highlighting the gap between laboratory-derived evolutionary potential and clinical prevalence is the R164N mutation. Despite exhibiting a strong resistance profile, with MIC values equal to or exceeding those of the clinically observed R164S variant [13] , R164N has only been detected experimentally and not in natural isolates [21]. This rarity presumably results from the requirement for two simultaneous nucleotide substitutions to achieve the R164N mutation, compared to only one nucleotide change necessary for R164S.

### Specialist Mutations: Antibiotic-Specific Adaptations

In contrast to generalist mutations that confer cross-resistance to multiple antibiotics, our analysis identified a distinct set of specialist mutations that provide resistance to only one β-lactam antibiotic (**Figure 2B, 2C**). Unlike the predictable localization of generalist mutations, specialist mutations showed broader structural distribution across the protein, appearing at diverse locations including both active site and peripheral regions. Analysis of the SML revealed significant differences in adaptive potential across tested β-lactams, with variation in both the abundance of specialist mutations and their unpredictable structural locations

For cefepime (FEP) and ceftriaxone (CFX), we identified no specialist mutations that met our significance criteria (log₂FoldChange > 3, p_adj_ < 0.05), suggesting limited evolutionary pathways for specific resistance to these antibiotics. For ampicillin (AMP), we observed only three specialist mutations that haven’t been observed before as a clinical isolate: R120S, S258D, T114G, none of which were located near the enzyme’s active site (**Figure 2B**). These mutations exhibited log₂FoldChange values only slightly above our threshold (log2FoldChange >3), suggesting they confer modest improvements in protein function rather than dramatic increases in resistance. This limited adaptive potential against ampicillin is consistent with TEM-1’s evolutionary history as an enzyme already optimized for ampicillin hydrolysis, leaving minimal room for further single-point mutation improvements. We also identified several strong adaptive mutations under aztreonam (AZT) selection, particularly E104R, A237G, and E240S. Notably, A237G has previously been observed in clinical isolates, typically in combination with other mutations. This variant is a component of TEM-22 (Q39K-E104K-A237G-G238S), as documented by Arlet et al. (1999) [22]. The other two positions (E104 and E240) are commonly substituted in clinical variants, typically E104K and E240K. The identification of alternative substitutions at these positions (E104R and E240S) demonstrates the broader mutational tolerance of these sites beyond what is typically observed in clinical isolates. This pattern suggests that while specific substitutions may dominate in clinical settings, the actual adaptive landscape at these positions is broader than what is typically observed in patient isolates.

Unlike generalist mutations that cluster predictably at three key functional positions, ceftazidime specialist mutations displayed remarkable structural diversity, appearing throughout the protein architecture. These mutations were predominantly located around the active site and concentrated on the Ω-loop but also appeared at the active site periphery and distant structural regions. This abundance suggests that ceftazidime presents unique selective pressure that can be overcome through multiple distinct molecular adaptations.

### The E166 Paradox: Catalytic Residue as an Evolutionary Hotspot

Among the specialist mutations identified in our analysis, those affecting position E166 were particularly intriguing and unexpected (**Figure 2B, 2C**). The E166 residue plays a direct and critical role in the β-lactam hydrolysis mechanism. β-lactam hydrolysis proceeds through a well-characterized two-step process: first, a nucleophilic attack by serine 70 on the β-lactam ring forms an acyl-enzyme adduct; subsequently, E166, with the assistance of an activated water molecule, performs the critical deacylation step that regenerates the active enzyme. This catalytic mechanism is conserved across β-lactamases, with positions S70 and E166 (following the Ambler nomenclature) being highly conserved among class A β -lactamases [23].

This discovery defies conventional mechanisms of hydrolysis and raises the question of whether TEM-1 possesses an alternative, secondary catalytic mechanism specifically capable of hydrolyzing ceftazidime but not other β-lactam antibiotics. Our data suggests that several substitutions at position E166 (E166P, E166G, E166F, E166Y, E166H, and E166C) can confer substantial resistance specifically to ceftazidime.

To verify the magnitude of resistance changes conferred by E166 mutations, we engineered four specific variants (E166G, E166H, E166D, and E166P) in the TEM-1 wild-type background and determined their Minimal Inhibitory Concentrations (MICs) against a panel of β-lactam antibiotics (**Table 1**). While wild-type TEM-1 confers robust protection against ampicillin (MIC: 30,000 µg/ml), the E166 variants displayed dramatically reduced MICs: 13.72 µg/ml (E166D), 4.57 µg/ml (E166G), 13.72 µg/ml (E166H), and 4.57 µg/ml (E166P), representing a three- to four-log decrease in resistance. These findings confirm the essential role of residue E166 in penicillin hydrolysis.

Remarkably, the same mutations led to increased resistance to ceftazidime, with MICs rising from 1.33 ± 0.0 µg/ml in wild-type TEM-1 to 4.00 ± 0.0 µg/ml in E166D, E166G, and E166H variants. Notably, E166P exhibited the most pronounced shift, with an MIC of 12.00 ± 0.0 µg/ml, despite showing no residual activity against ampicillin. In contrast, these mutations had no significant effect on resistance levels to other tested antibiotics, such as aztreonam, cefepime, cefotaxime, and ceftriaxone.

This finding raises a fundamental mechanistic question: if glutamic acid at position 166 is essential for the deacylation step, how do these mutants maintain catalytic activity against ceftazidime?

Biochemical evidence supporting this paradoxical substrate specificity comes from kinetic studies by Stojanoski et al. that examined E166Y in combination with P167G, another significant ceftazidime specialist identified in our screen. Kinetic analysis revealed that E166Y alone increased kcat/Km for ceftazidime by approximately 15-fold compared to wild-type TEM-1, while showing no enhancement for other antibiotics tested. When combined with P167G, the double mutant achieved a ∼35-fold increase in kcat/Km for ceftazidime and a modest 2.5-fold increase for cefotaxime. Most remarkably, a triple mutant (W165Y-E166Y-P167G) demonstrated a ∼400-fold increase in kcat/Km for ceftazidime [12]. Together with our MIC data, these kinetic findings **p**rovide compelling evidence that mutations in the Ω-loop region, particularly at position E166, can dramatically enhance ceftazidime hydrolysis through an alternative catalytic mechanism that remains to be fully characterized.

### Indirect β-Lactamase Activity Assay

To evaluate whether the TEM-1-E166P variant can hydrolyze ceftazidime, we performed an indirect β-lactamase activity assay by incubating four strains: non-resistant *E.coli* strain, TEM-1 wild-type, TEM-1-E166P variant, and TEM-1-R164N (positive control for ceftazidime hydrolysis) with various concentrations of ceftazidime and subsequently testing filtered supernatants for residual antibiotic activity using ceftazidime-sensitive *E.coli* strain (**Supplement Figure 3**). The rationale behind this assay was that strains incapable of degrading ceftazidime would leave the antibiotic concentration in the supernatant unchanged, resulting in a continued inhibitory effect on non-resistant *E.coli* growth. In contrast, strains that actively hydrolyze or sequester ceftazidime would reduce ceftazidime concentration during the initial incubation, allowing sensitive strain to grow when exposed to the treated supernatant. After 9 hours of primary incubation, we determined the initial minimal inhibitory concentration (MIC) for each tested strain. We observed a 2-fold increase in MIC of TEM-1 wild-type and an 8-fold increase in MIC of TEM-1-E166P (**Figure 3C**). Our positive control TEM-1-R164N had MIC 64-fold higher than the ceftazidime-susceptible *E.coli* strain. Next, cultures were centrifuged, and the resulting supernatants with concentrations lower or equal to 8 µg/mL were filtered to remove any remaining bacteria from β-lactamase-producing variants. These cell-free supernatants were then re-tested for residual antibiotic activity over a 16-hour period using the ceftazidime-sensitive *E.coli* strain by measuring endpoint MICs again. Initial incubation of cells with ceftazidime showed that supernatants from non-resistant *E.coli* and TEM-1 maintained MICs of 0.25 µg/mL, indicating negligible ceftazidime degradation. In contrast, *E.coli* cells exposed to E166P-treated supernatants showed a 4-fold increase in MIC (1 µg/mL) compared to supernatants obtained from non-resistant *E.coli* and TEM-1 *wild-type*. Notably, their growth was significantly impacted, showing very restricted growth, suggesting that residual ceftazidime remained in the supernatant. Actively hydrolyzing variant R164N-treated supernatants exhibited robust growth at all tested ceftazidime concentrations (≤ 8 µg/mL), confirming efficient hydrolysis (**Figure 3D**).

**Figure 3.**
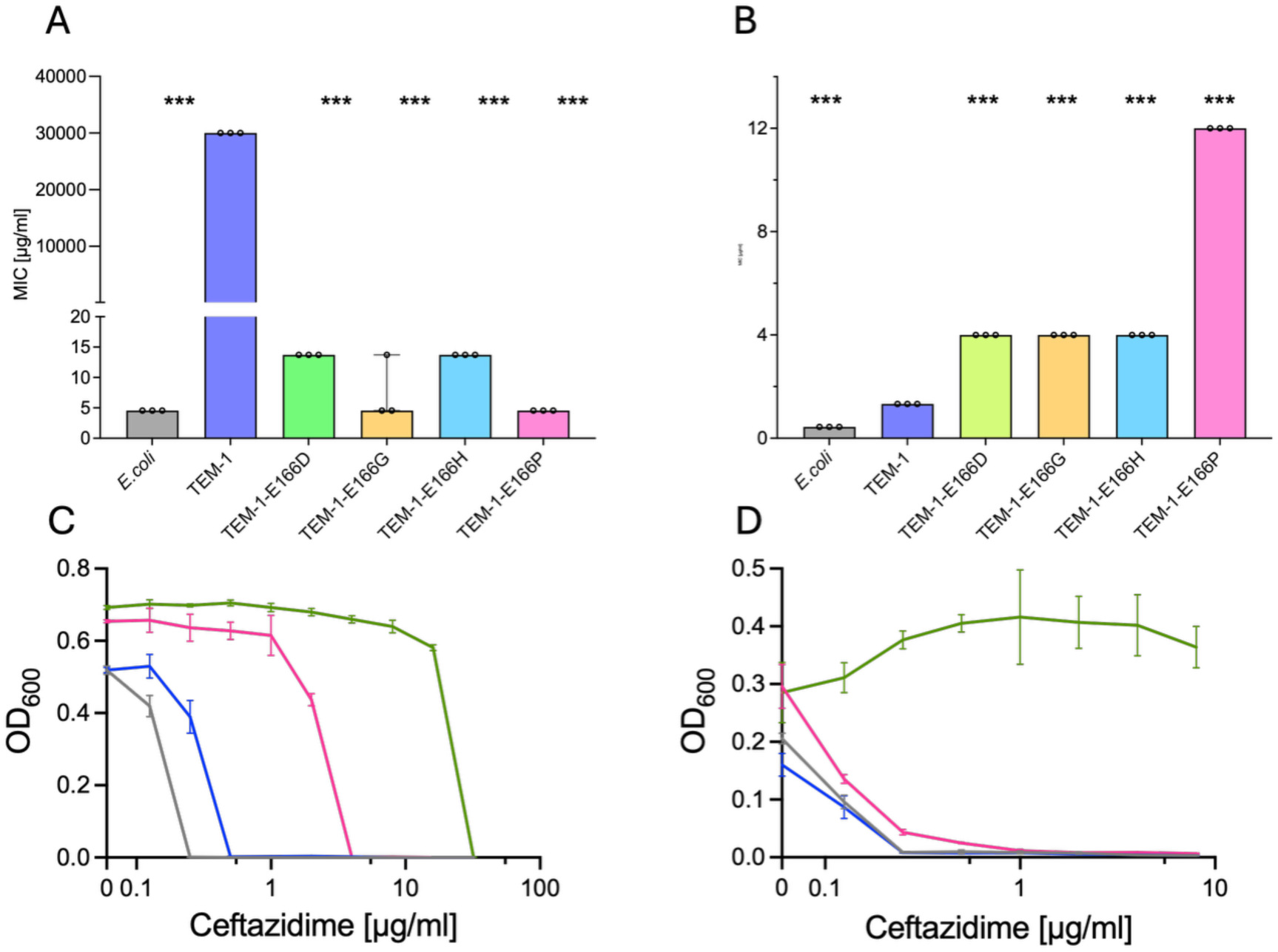
Resistance trade-off in TEM-1-E166P mutants with partial ceftazidime degradation detected by indirect β-lactamase assay. **(A, B)** Minimal inhibitory concentrations (MICs) against **(A)** ampicillin and **(B)** ceftazidime of the non-resistant *E. coli* NEB10β strain with and without *wild-type* TEM-1 or the indicated TEM-1 variants. Bars show the median ± 95 % confidence interval from three technical replicates (n = 3). Statistical differences versus wild-type were assessed with Mann–Whitney test (***p < 0.001). **(C)** Growth curves obtained during Indirect β-lactamase assay of the four strains during the 9 h primary incubation with a ceftazidime gradient. Colors: ceftazidime-sensitive *E. coli* control (black), TEM-1 wild-type (green), TEM-1-E166P (magenta), and TEM-1-R164N (blue). Points represent mean ± SD (n=3). **(D)** Growth curves of ceftazidime-sensitive *E.coli* obtained during Indirect β-lactamase assay strain after 16h of growth in filtered supernatants collected from the primary cultures in panel C. Colors correspond to the strain that produced each supernatant, as in panel C. Data are mean ± SD (n = 3).

These observations suggest that TEM-1-E166P grants limited ceftazidime tolerance, possibly through slow or incomplete hydrolysis or sequestration, rather than the robust degradation characteristic of true extended-spectrum TEM-1 variants.

### Molecular Dynamics Simulations Reveal Alternative Water Coordination Mechanisms in TEM-1-E166P

To investigate how the TEM-1-E166P variant might support ceftazidime hydrolysis despite lacking the essential general base E166, we performed Molecular Dynamics Simulations. We constructed acyl-enzyme complex models of wild-type TEM-1 and the E166P variant by superimposing the TEM-1 structure (PDB: 1BTL) to the ceftazidime-bound, deacylation-deficient KPC-2 structure (pdb: 6Z24). To mimic the covalent acyl-enzyme intermediate, we manually modified the ceftazidime molecule to form a covalent bond with the active site serine of TEM-1, thereby modeling the stable, pre-deacylation state of the complex. Each system was simulated in triplicate for 1 µs to examine water coordination patterns and potential alternative catalytic mechanisms.

Root-mean-square fluctuation (RMSF) analysis revealed no major differences in protein flexibility between wild-type TEM-1 and E166P, indicating that the resistance mechanism does not involve large-scale structural rearrangements (**Supplement Figure 4**). However, water positioning within the active site differed substantially between the variants. Both systems coordinated water molecules within 3.5 Å of the ceftazidime C6 carbon (the nucleophilic attack site) in 63% of simulation frames, but the spatial organization of these water molecules varied significantly (**Figure 4A, 4B**). In wild-type TEM-1, water molecules were optimally positioned for E166-mediated activation only 5.4% of the time, highlighting the dynamic nature of the catalytic process.

**Figure 4.**
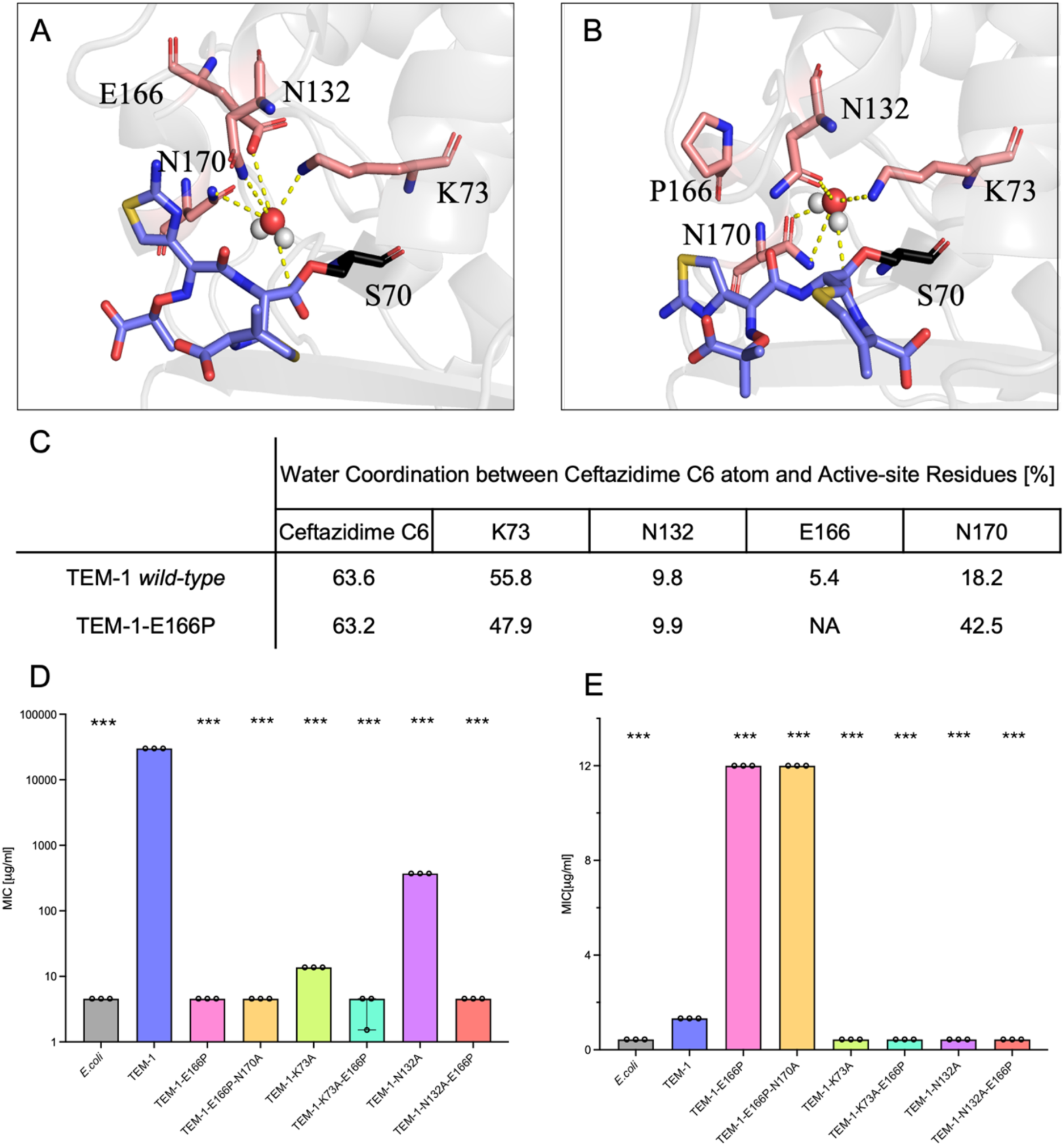
E166P Retains Water Coordination Mediated by N132 Catalytic Significance. **(A, B)**. Molecular Dynamic Simulation Snapshots Panel A, B represent a snapshot from Molecular Dynamic simulation of either TEM-1 (Panel A) or TEM-1-E166P (Panel B) of acyl-bound structure of Ceftazidime highlighting coordinated water molecules (represented as a sphere) by catalytically important residues shown as peach sticks: K73, N132, N170, and E166 (or P166). Hydrogen bonds are shown with yellow dashes for distances ≤ 3.5 Å. Ser70, to which the ligand is bound, is shown as a black stick. **(C)**. Water Molecule Coordination Table representing water molecules coordinated within 3.5 Å of Ceftazidime C6 carbon (nucleophilic attack site) and within 3.5 Å of K73, N132, N170, E166. Numbers are presented as centages from 3 simulation replicates of 1 µs each per protein. Panel **(D**, **E**) Minimal Inhibitory Concentrations against Ampicillin (D) and Ceftazidime (E) shown as a Median + 95% Confidence Interval for tested TEM-1 variants. Each strain was tested in 3 replicates. A two-tailed Mann–Whitney U test was performed to compare each variant against TEM-1 (*** p < 0.001)

Analysis of hydrogen bonding patterns revealed that K73 adopts different interaction modes between variants. In wild-type TEM-1, K73 frequently forms hydrogen bonds with E166, while in the E166P variant, K73 redirects to form hydrogen bonds with the oxygen of N132 **(Supplement Figure 6).** Intriguingly, the K73 side chain nitrogen frequently satisfied dual coordination criteria in both variants (55.5% in wild-type, 47.9% in E166P), while N170 showed enhanced water coordination in the E166P variant. Most notably, N132 maintained consistent water coordination behavior in both systems (∼9.8-9.9% of simulation time), though with different coordination modes between variants (**Figure 4C**).

To test the functional importance of these computationally identified residues, we constructed and characterized several mutant variants. The K73A mutation severely compromised all enzymatic activity regardless of the E166P background, confirming K73’s indispensable role in catalysis. More revealing was the behavior of N132A mutations. While TEM-1-N132A retained substantial ampicillin resistance (>90-fold higher than sensitive *E. coli*), it completely lost ceftazidime activity. Strikingly, the double mutant TEM-1-N132A-E166P lost resistance to both antibiotics, including the ceftazidime resistance normally conferred by E166P alone (**Figure 4D**). This finding aligns with extensive literature demonstrating N132’s critical role in cephalosporin hydrolysis. Previous studies by Judge et al. (2023) [24] and Jacob et al. (1990) [25] showed that N132 is essential for cephalosporin hydrolysis across class A β-lactamases, tolerating no amino acid substitutions and playing a crucial role in transition-state stabilization rather than ground-state binding. Our results extend these findings to TEM-1 and reveal that N132 plays an essential supporting role in the alternative catalytic mechanism employed by E166P.

To further probe the deacylation mechanism, we tested whether other active site residues could substitute for E166’s catalytic function. The variants E166P-E168A, E166P-N170A, and E166P-E168A-N170A all completely lost ampicillin hydrolysis activity (MIC: 4.57 µg/ml), confirming that neither E168 nor N170 can replace E166 in conventional penicillin hydrolysis. However, their ceftazidime resistance patterns were more complex. E166P-E168A showed only modest ceftazidime resistance (MIC: 4.00 µg/ml), similar to other catalytically impaired E166 variants. In contrast, both E166P-N170A and E166P-E168A-N170A maintained high ceftazidime resistance levels (MIC: 12.00 µg/ml), identical to E166P alone and representing a 9-fold increase over wild-type TEM-1 (**Supplement Figure 2.5**).

These results reveal that E166P employs an alternative catalytic mechanism for ceftazidime hydrolysis that does not rely on conventional general base-mediated water activation [26]. The E166P variant redirects K73 to form hydrogen bonds with N132, creating a distinct catalytic network that supports ceftazidime resistance through an unconventional deacylation pathway. While this K73-N132 interaction is essential for E166P’s activity, N132 appears to play a structural or stabilizing role rather than directly activating water molecules for deacylation. The mechanism likely involves enhanced positioning of catalytic water molecules and possibly facilitated water self-activation events within the reorganized active site architecture. Although this alternative pathway operates less efficiently than the conventional E166-mediated mechanism (consistent with the moderate ceftazidime resistance observed), it demonstrates that β-lactamases can evolve functionally competent resistance through non-canonical catalytic strategies. The absolute requirement for both K73 and N132 in this process highlights the precise structural organization needed to support this alternative resistance mechanism, representing a novel evolutionary strategy for acquiring activity against challenging substrates like ceftazidime.

## Discussion

Our study provides a comprehensive view of the mutational landscape of TEM-1 β-lactamase under selective pressure from diverse β-lactam antibiotics. Using a single mutant saturation mutagenesis library, we identified both generalist mutations that confer broad-spectrum resistance and specialist mutations that drive antibiotic-specific adaptation. These findings underscore the evolutionary flexibility of TEM-1, including positions previously assumed to be mechanistically constrained.

Consistent with prior knowledge, we found that TEM-1 is highly optimized for penicillin hydrolysis, as reflected by the limited number and impact of resistance-conferring mutations under ampicillin selection [8]. Only a small fraction of generalist mutations resides within locations important for protein activity. Mutations localized on R164 position are important for omega loop flexibility, which allows accommodation of bulky β-lactam antibiotics common in higher generation cephalosporins or potentially enables better positioning of E166 for the deacylation step [7, 27]. Two other positions, E240 and G238, are directly present within the active site and can directly impact drug binding. G238S, commonly observed in clinical isolates, is speculated to improve antibiotic binding affinity [18, 28, 29].

Specialist mutations showed a wider range of unique adaptations. Ceftazidime specialist mutations presented a strikingly different adaptive profile, yielding the greatest number and diversity of beneficial mutations. Many of these mutations clustered around the Ω-loop, suggesting that even minor alterations in this region can significantly affect substrate binding and catalysis [12, 15, 21]. This supports the notion that ceftazidime imposes a unique structural challenge on TEM-1, one that the enzyme can overcome through multiple mutation-driven solutions [12, 30–32].

A specialist mutation that merited further investigation was the E166P variant observed under ceftazidime selection. While previous studies have documented E166 mutations arising under ceftazidime pressure [12, 24], none have explored the mechanistic basis of how E166P could retain hydrolytic activity. This variant displayed a remarkable and paradoxical phenotype: a complete loss of ampicillin resistance coupled with enhanced resistance to ceftazidime. This striking phenotypic trade-off challenged the conventional understanding of TEM-1 catalysis and prompted us to investigate whether the generally accepted catalytic mechanism, in which E166 serves as the general base to activate the water molecule required for deacylation, is replaced by an alternative mechanism specifically for ceftazidime hydrolysis.

Our indirect β-lactamase activity assay supported this hypothesis. Supernatants from cultures expressing E166P displayed increased MICs against the ceftazidime-sensitive *E.coli* strain, suggesting that ceftazidime concentration decreased despite the loss of a functional general base, but the restricted growth of *E.coli* nonresistant strain, suggested that residual ceftazidime remained in the supernatant. This result suggests that E166P may either form a stable acyl-enzyme adduct that reduces free ceftazidime levels through sequestration or facilitate an alternative slow hydrolysis mechanism.

Molecular dynamics simulations revealed the structural basis for this alternative catalytic capability. While both wild-type TEM-1 and E166P coordinate water molecules near the C6 carbon of ceftazidime with similar frequency (63% of simulation frames), the critical difference lies in the organization of the active site hydrogen bonding network. Our most significant discovery was that K73 adopts fundamentally different interaction modes between variants: in wild-type TEM-1, K73 forms hydrogen bonds with E166, while in the E166P variant, K73 redirects to form hydrogen bonds with N132, creating an entirely reorganized substrate recognition and positioning system.

This K73-N132 network proves essential for E166P’s ceftazidime-specific activity. Functional validation through mutagenesis confirmed that while K73 remains absolutely critical in both variants, N132 emerges as the key enabler of the alternative mechanism. The complete loss of ceftazidime resistance in the TEM-1-N132A-E166P double mutant demonstrates that N132 serves a dual role: maintaining transition-state stabilization in wild-type enzyme while functioning as a critical substrate positioning element in the E166P variant. Without N132, the reorganized active site cannot properly bind or position ceftazidime for hydrolysis, explaining the total elimination of resistance.

The enhanced water coordination activity of N170 in the E166P variant, coupled with the finding that N170 removal does not impair E166P’s ceftazidime activity, further supports the primacy of the K73-N132 axis in the alternative mechanism. This selectivity indicates that the E166P variant has evolved a highly specific substrate recognition system that bypasses the conventional E166-mediated pathway.

Our findings align with previous observations that ceftazidime presents unique challenges to TEM-1 β-lactamases due to its bulky side chain and structural complexity [12]. The transient nature of optimal catalytic water positioning in wild-type TEM-1 (observed in only 5.4% of simulation time) suggests inherent inefficiency in ceftazidime hydrolysis through the traditional E166-mediated mechanism. The E166P variant appears to have circumvented this limitation by developing an alternative substrate engagement strategy that, while less efficient than conventional mechanisms, provides sufficient catalytic capability for meaningful resistance.

Our results are consistent with the broader pattern of ceftazidime resistance evolution across β-lactamase families. Judge et al. demonstrated that E166, P167, and N170 mutations in CTX-M14 specifically enhance ceftazidime resistance by increasing omega loop flexibility [24]. The clinical isolation of TEM-193 (E166G) with ten additional compensatory mutations further illustrates the structural constraints and evolutionary pressure associated with disrupting this critical catalytic residue[33].

The specialist nature of the E166P adaptation contrasts sharply with broad-spectrum resistance mutations that typically emerge under multi-drug selective pressure. While the E166P variant shows remarkable specificity for ceftazidime resistance with minimal impact on ampicillin activity, this narrow spectrum adaptation would likely be disadvantageous in environments with diverse β-lactam exposure. However, in clinical settings where ceftazidime dominates therapeutic protocols, such specialist variants could emerge and persist.

Importantly, our findings demonstrate that while single point mutations provide valuable insights into evolutionary possibilities, TEM-1 typically requires multiple mutations to become an extended-spectrum β-lactamase capable of efficiently hydrolyzing later-generation cephalosporins and monobactams [31, 32, 34]. This requirement for combinatorial mutations explains why clinical isolates commonly harbor between 2-5 mutations rather than single substitutions [7, 33]. The synergistic effects of multiple mutations likely overcome the individual limitations of single-residue changes, enabling the robust resistance profiles observed in clinical extended-spectrum β-lactamases[21, 35].

Our work demonstrates that even highly conserved catalytic residues can serve as evolutionary innovation points under appropriate selective pressure. The E166P variant exemplifies how structural constraints can be circumvented through alternative biochemical mechanisms involving reorganization of substrate recognition networks rather than modification of catalytic chemistry itself. This mechanistic flexibility underscores the ongoing challenge of combating antibiotic resistance and the need for comprehensive approaches that account for both functional and structural diversity in resistance mechanisms.

## Acknowledgements

This work is supported by the UT Southwestern Endowed Scholars Program (E.T. and M.M.L), the Welch Foundation I-2082-20240404 (E.T.), the Alfred P. Sloan Foundation G-2024-22449 (M.M.L.), and the NIH NIGMS R01GM125748 (E.T.) and R35GM150897-01 (M.M.L.).

## Methods

### Growth Media and Strains

Bacteria cells were grown at 37°C in Luria Broth media (L24040, RPI) supplemented with 12µg/ml Tetracycline hydrochloride (NA-0254, Combi-Blocks) except *E. coli* NEB10β cells that do not carry the plasmid. All *E. coli* strains NEB10-beta *wild-type* are DH10B derivative.

### Antibiotic Compounds

The antibiotics used in the experiments were: Ampicillin sodium salt (A0166, Sigma-Aldrich), Cefotaxime sodium (SS-7557, Combi-Blocks), Azactam/Aztreonam for Injection (NDC 0003-2560-16, Bristol-Myers Squibb), Cefepime for Injection (NDC 60505-6146-0, Apotex Corp.), Ceftriaxone for Injection (NDC 0409-7332-01, Hospira), Ceftazidime for Injection (NDC 44567-235-25, WG Critical Care), and Tetracycline hydrochloride (NA-0254, Combi-Blocks). All antibiotics, except for tetracycline hydrochloride, were dissolved in Molecular Biology Grade Water and filtered through a 0.2 µm syringe filter. Tetracycline hydrochloride was dissolved in 70% ethanol and similarly filtered through a 0.2 µm syringe filter.

### Transformation and pooling of the saturation mutagenesis library of TEM-1 β-lactamase

The construction of the TEM-1 β-lactamase saturation mutagenesis library was carried out and described by Michael Stiffler[8]. The saturation mutagenesis library was constructed on the low-copy plasmid pBR322, which contains a constitutively active promoter. Library were divided into ten sub-groups based on amino acid positions (26–51, 52–78, 79–104, 105–132, 133–156, 157–183, 184–209, 210–236, 237–264, 265–290), covering the entire mature protein sequence. Each sub-group (4 µl) was independently transformed in triplicate into 20 µl of *E. coli* NEB10B electrocompetent cells via electroporation. After a 1-hour incubation at 37°C with shaking at 230 rpm, 10 µl of the culture was taken to assess transformation efficiency. Each transformation yielded greater than 10⁶ CFU/ml. The remaining culture was transferred to 50 ml LB medium supplemented with 12 µg/ml tetracycline and incubated overnight at 37°C, 230 rpm. The following day, the OD600 of each sub-library was measured and adjusted to 2.0. Subsequently, all sub-libraries were pooled together in equal OD600 ratios to generate three independent Saturation Mutagenesis Libraries of TEM-1. The pooled sub-libraries were then aliquoted into independent glycerol stocks and stored at −80°C for further experiments.

### Selection Experiment of SML TEM-1

Three biological replicas of saturation mutagenesis libraries were thawed from the glycerol stocks and transferred to Erlenmeyer flasks containing 100 mL of LB media supplemented with 12 %g/mL Tetracycline. TEM-1 wild-type was streaked from a colony on an LB agar plate supplemented by Tetracycline 12 %g/mL and inoculated in 5 mL of previously described media. All cultures were grown overnight at 37°C with shaking at 230 rpm. Next morning, all overnight cultures were diluted 1:5 into 10 mL of media as described above for adaptation. The cells were incubated for 3 hours at 37°C with shaking at 230 rpm. After 3 hours, the OD600 of each culture was measured and diluted to 0.02 in a total volume of 10 mL of the media described above. Next, 100 %L of each diluted culture was transferred in duplicate to a pre-prepared 96-well plate, which contained 100 %L per well of media with the following final antibiotic concentrations after adding the cells:

- Ampicillin: 16,384, 8,192, 4,096, 2,048 %g/mL
- Ceftazidime: 16, 8, 4, 2, 1, 0.5, 0.25 %g/mL
- Aztreonam: 8, 4, 2, 1, 0.5, 0.25 %g/mL
- Cefepime: 8, 4, 2, 1, 0.5 %g/mL
- Ceftriaxone: 2, 1, 0.5, 0.25, 0.125, 0.0625 %g/mL
- Cefotaxime: 2, 1, 0.5, 0.25, 0.125, 0.0625 %g/mL

Additionally, all drug conditions included drug-free media as a control. The cells were incubated, and growth was monitored during the selection experiment for 18 hours, with shaking every 2 minutes for 20 seconds at 37°C. Next, Saturation Mutagenesis Libraries (SMLs) with drug free media and the one that exhibited growth at higher antibiotic concentrations than the TEM-1 wild-type control were selected for sequencing. Those wells were transferred to 20 mL of β-lactam antibiotic-free media and incubated for 3 hours at 37°C with shaking to increase the density of resistant cells and enhance plasmid concentration for sequencing. The selected SML wells for sequencing corresponded to the following concentrations:

- Ampicillin: 4,096 %g/mL,
- Ceftazidime: 1 %g/mL,
- Cefotaxime: 0.25 %g/mL,
- Ceftriaxone: 0.25 %g/mL,
- Aztreonam: 0.5 %g/mL,
- Cefepime: 1 %g/mL
- No drug

Next, those cells were centrifuged for 5 min at speed 6000rpm. Supernatant was discarded and pellet was store at -80°C.

### NGS Sample preparation

Each pellet obtained during the Selection Experiment of SML TEM-1 was thawed, and plasmid DNA was purified following the manufacturer’s protocol for the NucleoSpin Plasmid Kit (Macherey-Nagel). Next, the sequencing region was amplified using the following primer set: forward :(5’-TCAAATATGTATCCGCTCATGAGAC-3’) reverse (5’ AAAGGATCTTCACCTAGGTCCTT-3’).

The samples were heated to 98°C for 3 minutes, followed by 20 cycles of: 98°C for 15 seconds, 64°C for 20 seconds, and 72°C for 1 minute. A final extension step was performed at 72°C for 3 minutes. The amplified DNA was purified by gel extraction using the NucleoSpin Gel and PCR Clean-Up Kit (Macherey-Nagel). DNA quality was assessed using a spectrophotometer by measuring the 260/280 nm and 260/230 nm ratios, while DNA concentration was determined using a Qubit 3 Fluorometer (Thermo Scientific). The purified DNA products were then sent for 400 bp Illumina sequencing.

### NGS Data analysis

Paired-end 2 × 150 bp reads were aligned to the reference plasmid and translated in silico. Any read that showed more than a single amino acid change was discarded, and all reads with a single amino acid change assumed to not have any other mutations. Reads were collapsed by amino acid change, yielding an integer count for each variant across all replicate/drug combinations. This process generated two matrices per drug: mut_fit [m × n]: counts of each mutant (rows) across n = 3 biological replicates (columns) and wt_fit [w × n]: counts of synonymous WT codons across the same replicates. Read counts were normalized to frequencies within each replicate. To handle zero counts in log-transformation, a pseudocount of 0.1 was added prior to normalization. For mutant i in replicate r, fitness of each mutant was defined as enrichment as log fold change of frequencies with respect to the untreated culture:

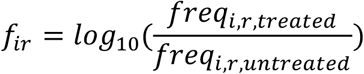

All statistical analyses were conducted in Python 3.11 using SciPy 1.11 and statsmodels 0.14. For each mutant, the null hypothesis “fitness equals the median fitness of synonymous wild-type” was tested using a paired Student’s t-test:

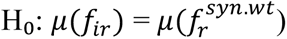

Here, each pair 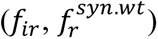 was taken from the same culture replicate r. The test was implemented using scipy.stats.ttest_rel. P-values were adjusted for false discovery rate (FDR) using the Benjamini–Hochberg procedure (α = 0.05) via statsmodels.stats.multitest.multipletests. Mutants with FDR-adjusted p < 0.05 were considered to have significantly different fitness from median synonymous WT.

### Minimal Inhibitory Concentrations

Minimum inhibitory concentration (MIC) was determined from OD₆₀₀ measurements taken after 20 h of growth in increasing antibiotic concentrations. For each strain and antibiotic, MIC was defined as the lowest concentration at which the background-corrected OD (OD₆₀₀ minus the plate’s median background) fell below 0.01. MIC values were calculated independently for each biological replicate, each of which comprised three technical replicates. Statistics are reported as the median MIC with a 95 % confidence interval estimated by bootstrap resampling. Pairwise comparisons of MIC distributions between strains were made using a two-tailed Mann–Whitney U test. Significance is denoted as * p < 0.05, ** p < 0.01, and *** p < 0.001.

### Indirect β-Lactamase Activity Assay

To assess the ability of TEM-1 variants to hydrolyze ceftazidime, we performed an indirect β-lactamase activity assay using four *E. coli* strains: NEB10B (non-resistant control), NEB10B carrying a plasmid encoding TEM-1 wild-type, NEB10B expressing TEM-1-E166P and NEB10B expressing TEM-1-R164N (ceftazidime-resistant variant).Overnight cultures were inoculated into 5 mL of Luria Broth medium. Strains carrying plasmids were grown in Luria Broth supplemented with tetracycline at a final concentration of 12 µg/mL, while the NEB10B control strain was grown in plain Luria Broth. All cultures were incubated overnight at 37°C with shaking. The following day, each overnight culture was diluted to an OD₆₀₀ of 0.001 in 3 mL of fresh plain LB medium containing one of the following concentrations of ceftazidime: 32, 16, 8, 4, 2, 1, 0.5, 0.25, 0.125, or 0 µg/mL. Cultures were then incubated for 9 hours at 37°C with shaking. As a control, tubes containing ceftazidime, but no bacterial cells were processed in parallel. All experimental conditions were performed in triplicate. After the 9-hour incubation, 200 µL of each culture was transferred to a 96-well plate to assess MIC using a BioTek spectrophotometer. For samples with ceftazidime concentrations equal to or below 8 µg/mL, cultures were centrifuged at 6000 rpm for 3 minutes. The supernatant from each was then filtered using 0.22 µm Spin-X centrifuge tube filters by spinning at 11,000 × g for 1 minute. A volume of 200 µL of the filtered supernatant, corresponding to each specific condition, strain, and replicate, was transferred to new wells in a 96-well plate. To evaluate residual antibiotic activity, 5 µL of an NEB10B overnight culture was added to each well containing the filtered supernatant. This inoculum was diluted in 10× Luria Broth medium to provide additional nutrients and adjusted to achieve a final OD₆₀₀ of 0.001 in each well. The plates were incubated for 16 hours at 37°C, 400 rpm, and 80% humidity. MICs were determined by evaluating the inhibition of NEB10B growth at the endpoint in each condition.

### Molecular Dynamics Simulations

The molecular dynamics system was prepared using the CHARMM-GUI Solution Builder platform. The protein structure was placed in a rectangular simulation box with a 1 nm edge distance and solvated using TIP3P water molecules. The system was neutralized and ionized by adding K⁺ and Cl⁻ ions to achieve an isotonic concentration of 0.15 M, mimicking physiological conditions. The simulation pH was set to 7.0. The generated system was first energy minimized for 5000 steps, using a convergence tolerance of 100.0 kJ/mol/nm². Initial velocities were generated at 300 K using a random seed. Electrostatics were treated with the Particle Mesh Ewald (PME) method with an Ewald tolerance of 0.0005 and van der Waals (vdW) interactions were handled using the force-switch method, with switch-on and switch-off distances set to 1.0 nm and 1.2 nm, respectively. Following minimization, a two-stage equilibration protocol was performed. The first equilibration was conducted for 5 ns (5,000,000 steps at 2 fs timestep) under NVT conditions with constant volume and temperature maintained at 300 K using Langevin dynamics with a friction coefficient of 1 ps⁻¹. During this phase, positional restraints were applied to both protein backbone and side chains with a force constant of 4000 kJ/mol/nm². The second equilibration was run for an additional 5 ns (5,000,000 steps) under NPT conditions, with a Monte Carlo barostat applied at 1 bar pressure using isotropic pressure coupling. The production simulation was then carried out for 1 µs (50,000,000 steps) using the same NPT settings, but with all positional restraints removed. Coordinates were recorded every 50,000 steps, and simulation outputs were written every 1,000 steps. Each system was simulated in triplicate to ensure reproducibility and statistical significance of the results.

### Water Analysis

We quantified bridging water molecules between ceftazidime acyl-carbon (C6) and active-site residues using MDTraj. We corrected periodic-boundary conditions to maintain ligand and residue integrity within the primary unit cell. Using md.compute_neighbors with a 0.35 nm cutoff (3.5 Å), we identified water-oxygen atoms near target atoms (C6; Lys73-NZ; Asp132-OD1/ND2; Glu166-OE1/OE2; Asp170-OD1/ND2). A water molecule was classified as bridging if simultaneously located ≤ 0.35 nm from C6 and an active-site residue atom. We calculated bridging waters per frame as a percentage of total frames. To validate, we randomly sampled frames and recomputed pairwise distances, confirming each identified water met the ≤ 0.35 nm threshold.

## Supplementary Figures

**Supplement Figure 1.**
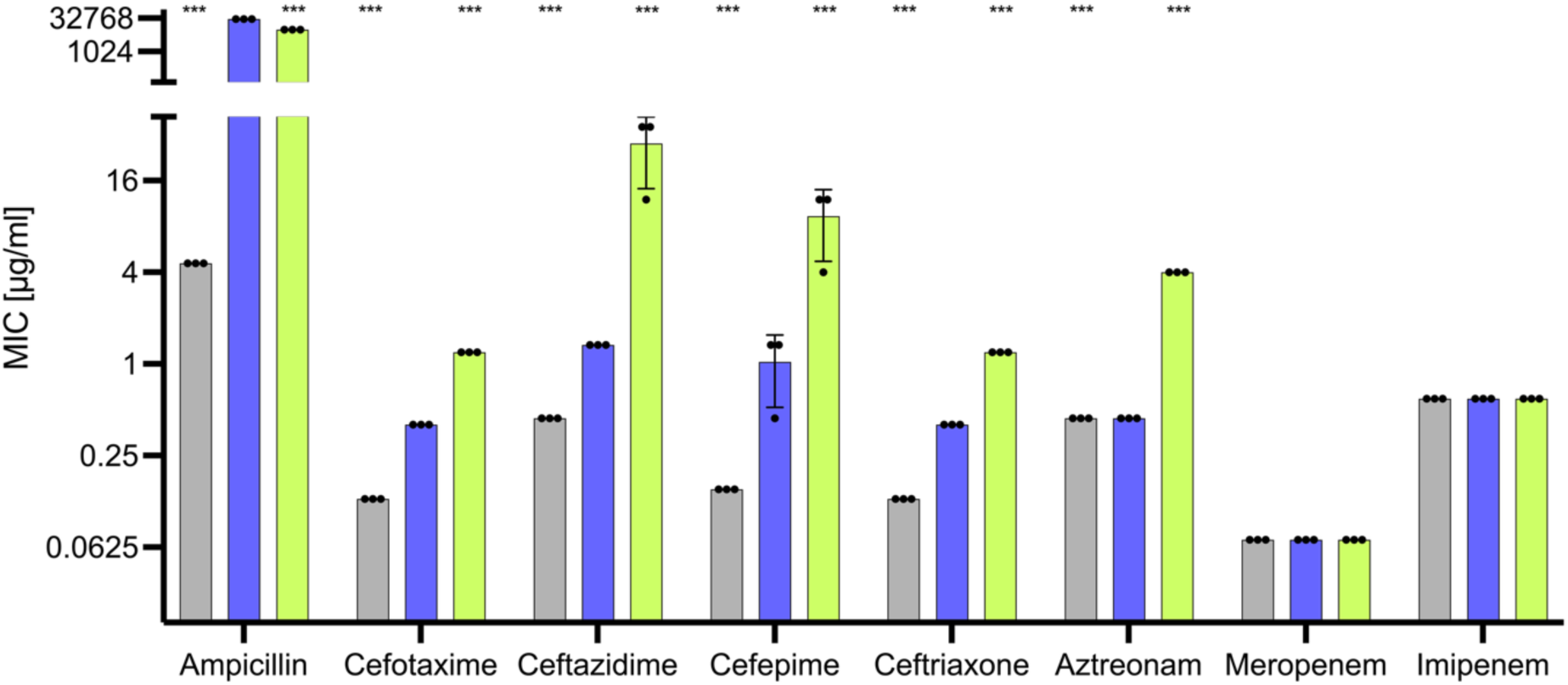
Resistance Profiles of TEM-1 Saturation Mutagenesis Library Against β-Lactam Antibiotics. Bar plot represents Minimal Inhibitory Concentrations of the non-resistant *E. coli* NEB10β strain (gray) and NEB10β strains producing *wild-type* TEM-1 (purple) and a pool of the Saturation Mutagenesis Library (light green) against representatives from four major β-lactam classes: penicillins (ampicillin), cephalosporins (cefotaxime, ceftriaxone, ceftazidime, cefepime), monobactams (aztreonam), and carbapenems (meropenem, imipenem). Bars show the median ± 95% confidence interval from three technical replicates (n = 3). Statistical differences versus wild-type were assessed with Mann–Whitney test (***p < 0.001).

**Supplement Figure 2.**
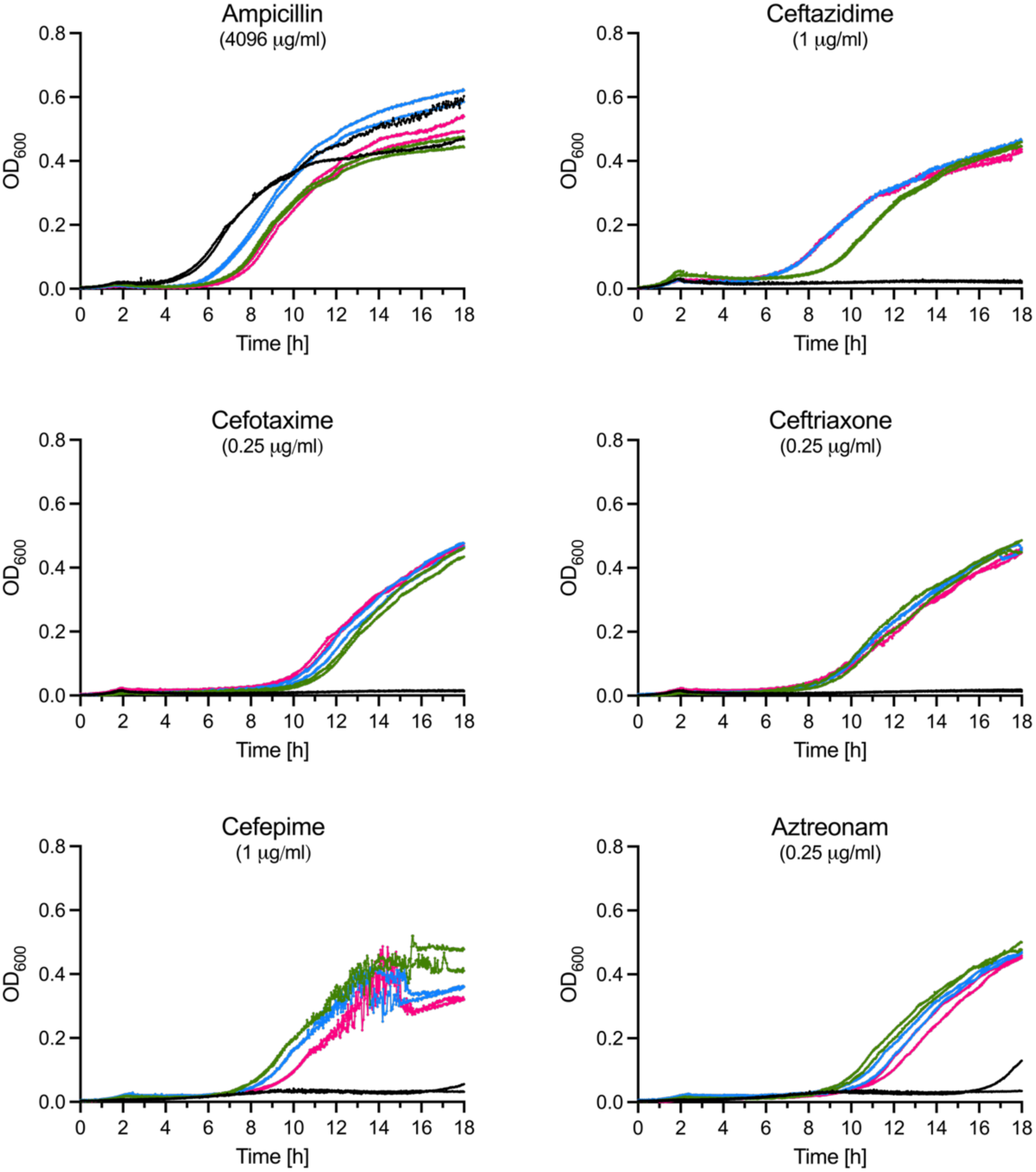
Selection of Antibiotic-Resistant Variants from the TEM-1 Single Mutant Library. Growth curves of *E. coli* harboring TEM-1 plasmid (black, control) or Saturation Mutagenesis Library (blue, magenta, green) during 18-hour antibiotic exposure in LB media at the concentrations selected for sequencing. Three biological replicates with two technical replicates were performed.

**Supplement Table 1.** Minimal inhibitory concentrations (MICs) of *E. coli* NEB10β and NEB10β strains expressing TEM-1 β-lactamase or its variants against β-lactam antibiotics.

| Strain | Minimal Inhibitory Concentrations [μg/ml] | | | | | |
| --- | --- | --- | --- | --- | --- | --- |
|  | AMP | CAZ | CTX | CFX | FEP | AZT |
| *E. coli* | 4.57 | 0.44 | 0.13 | 0.13 | 0.15 | 0.44 |
| TEM-1 *wild-type* | 30000 | 1.33 | 0.4 | 0.4 | 1.33 | 0.44 |
| TEM-1-E166D | 13.72 | 4 | 0.4 | 0.4 | 0.44 | 0.44 |
| TEM-1-E166G | 4.57 | 4 | 0.4 | 0.4 | 0.44 | 0.44 |
| TEM-1-E166H | 13.72 | 4 | 0.4 | 0.4 | 0.44 | 0.44 |
| TEM-1-E166P | 4.57 | 12 | 0.4 | 0.4 | 1.33 | 0.44 |

**Supplement Figure 3.**
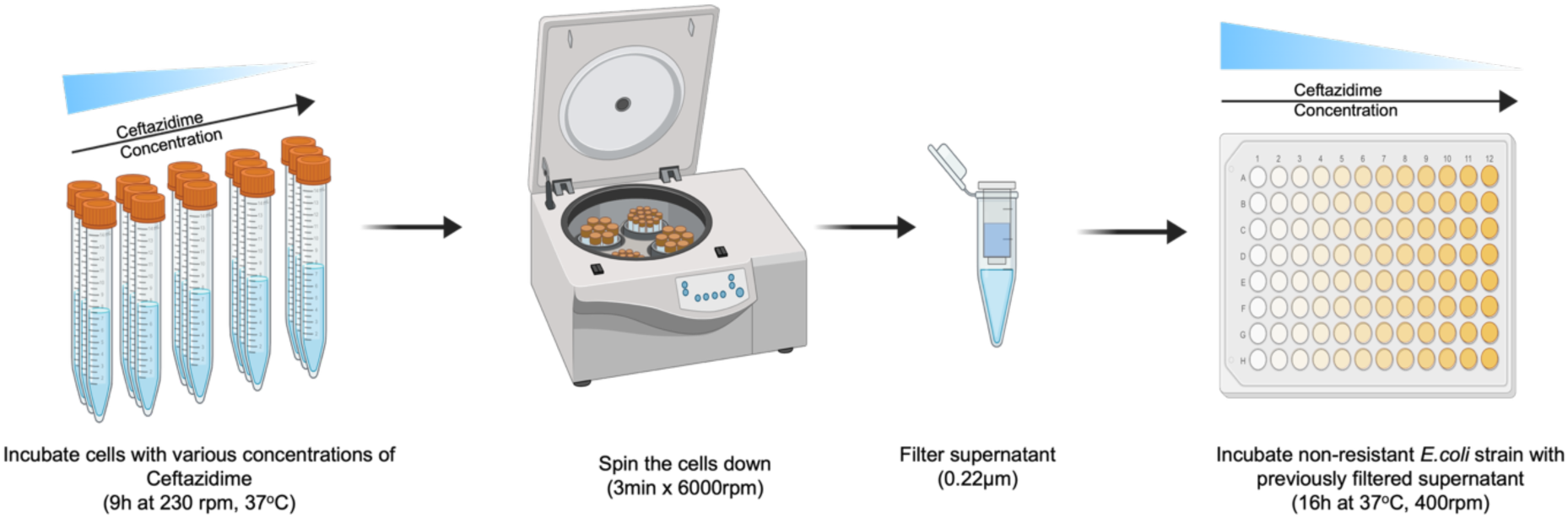
Indirect β-lactamase assay workflow. Strains were cultured for 9 h in a ceftazidime gradient, centrifuged, and the supernatant was passed through a 0.22 µm filter. The cell-free supernatants were then transferred to fresh 96-well plates, inoculated with the ceftazidime-sensitive *E. coli* strain, and incubated for 16 h. MIC measurement was performed at the end of each incubation.

**Supplement Figure 4.**
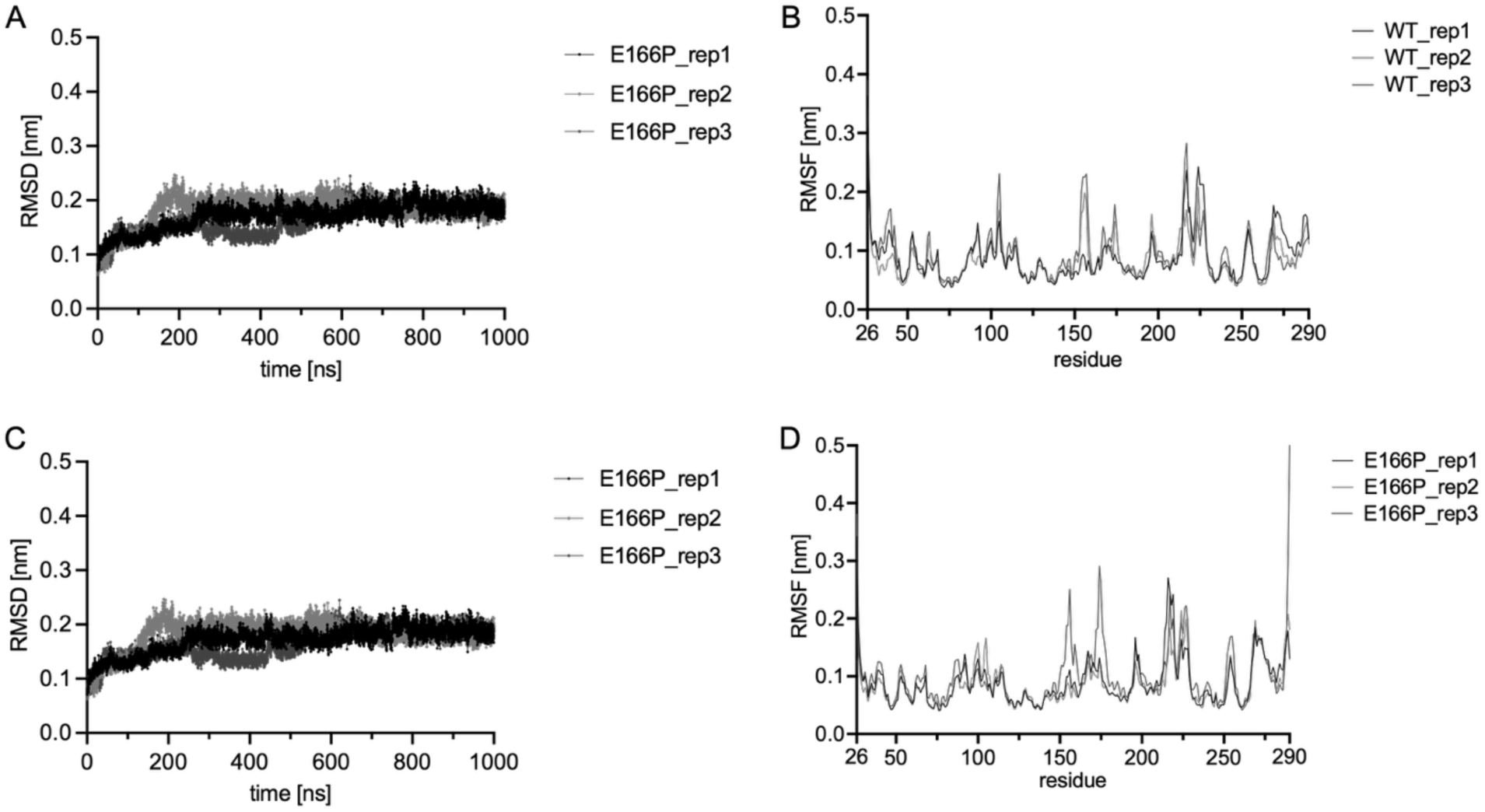
Molecular Dynamics Simulation Analysis of TEM-1 and TEM-1-E166P Variants. **(A, B)** Show Root Mean Square Deviation (RMSD) for TEM-1 (A) and TEM-1-E166P (B) shown for each replicate. **(C, D)** Show Root Mean Square Fluctuation (RMSF) for TEM-1(C) and TEM-1-E166P (D), shown as lines for each replicate obtained for Cα atoms. All molecular dynamics simulations were performed for 1 µs in triplicate

**Supplement Figure 5.**
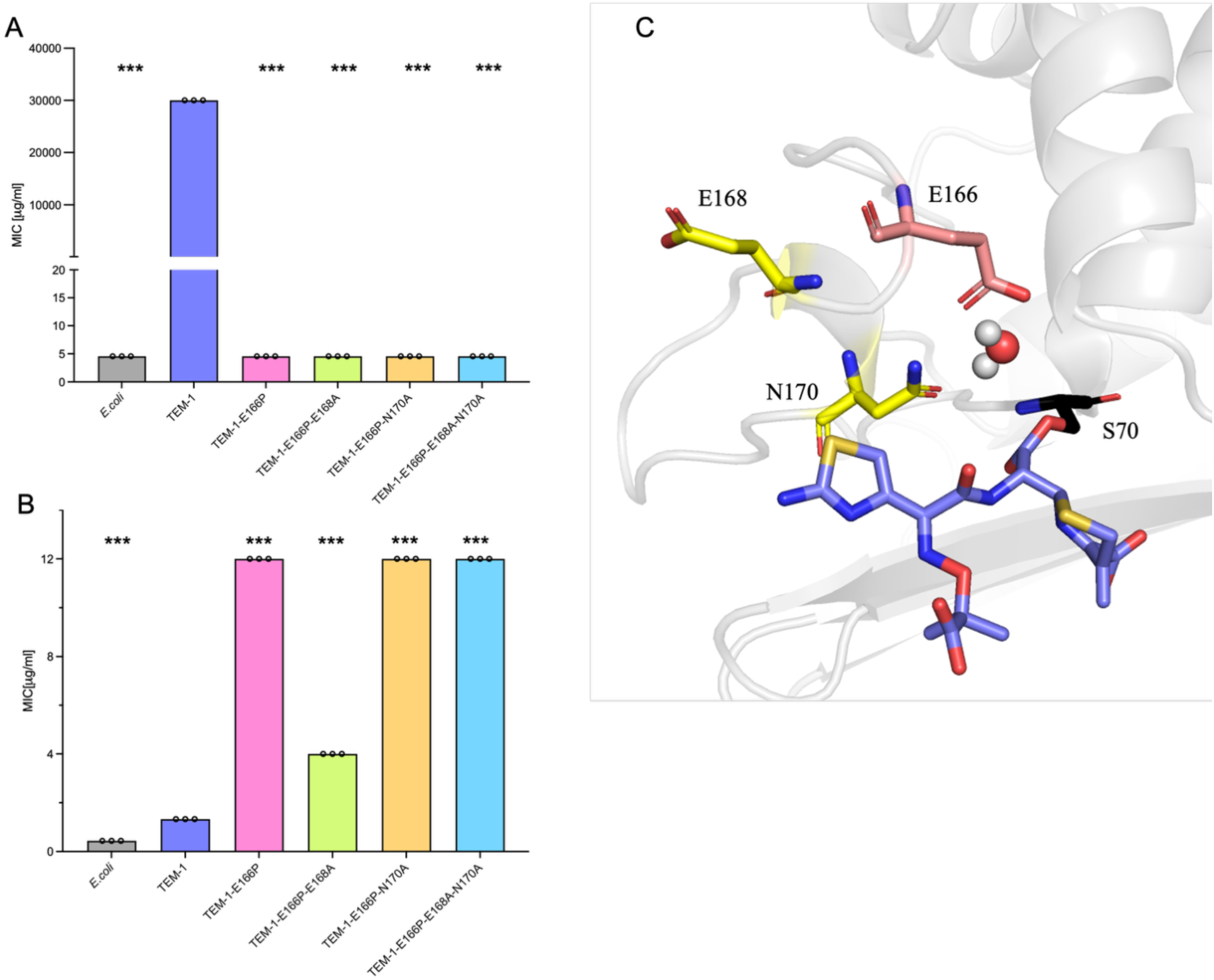
E168 and N170 Do Not Mediate Deacylation of the Ceftazidime Acyl-Enzyme in the TEM-1–E166P Variant. **(A, B)** Minimal inhibitory concentrations (MICs) of the non-resistant *E. coli* NEB10β strain and NEB10β strains producing *wild-type* TEM-1 or the indicated TEM-1 variants. Bars show the median ± 95 % confidence interval from three technical replicates (n = 3). Statistical differences versus wild-type were assessed with Mann–Whitney test (***p < 0.001). Panel A: MICs for ampicillin. Panel B: MICs for ceftazidime. **(C)** Acyl-enzyme model of wild-type TEM-1 (PDB 1BTL) with covalently bound ceftazidime (purple). The catalytic Ser70 that forms the acyl bond is shown in black; the crystallographic water poised for nucleophilic attack is shown as a sphere. The general base Glu166 is shown as peach sticks. Tested residues Glu168 and Asn170 are highlighted in yellow.

**Supplement Figure 6.**
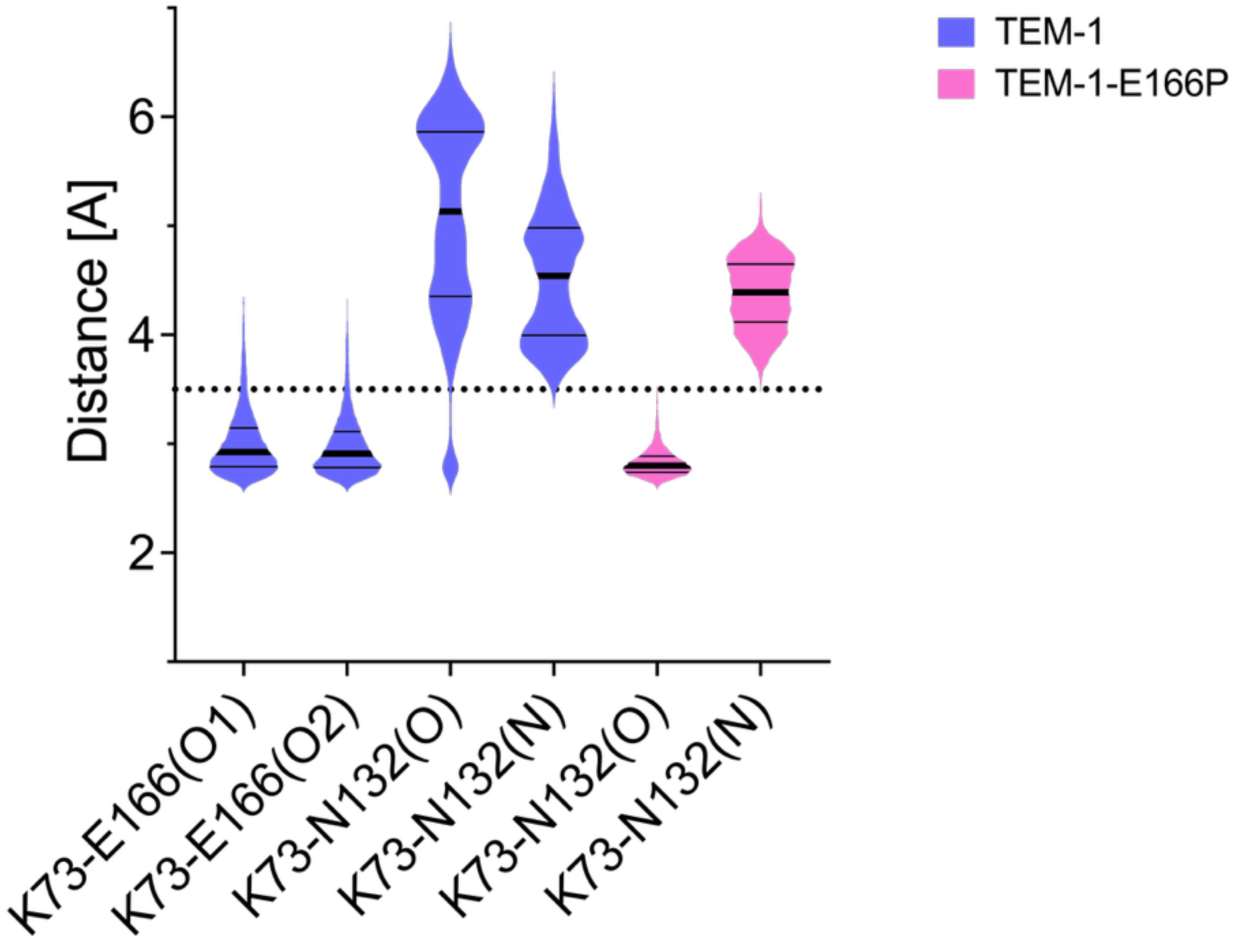
K73 Side Chain Interactions with E166 and N132 in TEM-1 and TEM-1-E166P Variants. Violin plots show the distribution of distances (in Å) between the side chain of residue K73 and either E166 (O1, O2) or N132 (O, N) observed in molecular dynamics simulations of TEM-1 (blue) and TEM-1-E166P (pink), each performed in triplicate. In TEM-1, K73 maintains close interaction with E166 (O1/O2), consistent with a catalytic hydrogen bond. In contrast, the TEM-1E166P forms a stable hydrogen bond with the side chain oxygen of N132. This shift suggests a compensatory interaction in TEM-1-E166P that may underlie its altered catalytic mechanism. The dotted line at 3.5 Å marks the conventional hydrogen bond distance threshold

